# Endogenous inflammatory mediators produced by injury activate TRPV1 and TRPA1 nociceptors to induce sexually dimorphic cold pain that is dependent on TRPM8 and GFRα3

**DOI:** 10.1101/2023.01.23.525238

**Authors:** Chenyu Yang, Shanni Yamaki, Tyler Jung, Brian Kim, Ryan Huyhn, David D. McKemy

## Abstract

The detection of environmental temperatures is critical for survival, yet inappropriate responses to thermal stimuli can have a negative impact on overall health. The physiological effect of cold is distinct among somatosensory modalities in that it is soothing and analgesic, but also agonizing in the context of tissue damage. Inflammatory mediators produced during injury activate nociceptors to release neuropeptides, such as CGRP and substance P, inducing neurogenic inflammation which further exasperates pain. Many inflammatory mediators induce sensitization to heat and mechanical stimuli but, conversely, inhibit cold responsiveness, and the identity of molecules inducing cold pain peripherally is enigmatic, as are the cellular and molecular mechanisms altering cold sensitivity. Here, we asked if inflammatory mediators that induce neurogenic inflammation via the nociceptive ion channels TRPV1 and TRPA1 lead to cold pain in mice. Specifically, we tested cold sensitivity in mice after intraplantar injection of lysophosphatidic acid (LPA) or 4-hydroxy-2-nonenal (4HNE), finding each induces cold pain that is dependent on the cold-gated channel TRPM8. Inhibition of either CGRP, substance P, or toll-like receptor 4 (TLR4) signaling attenuates this phenotype, and each neuropeptide produces TRPM8-dependent cold pain directly. Further, the inhibition of CGRP or TLR4 signaling alleviates cold allodynia differentially by sex. Lastly, we find that cold pain induced by inflammatory mediators and neuropeptides requires the neurotrophin artemin and its receptor GFRα3. These results demonstrate that tissue damage alters cold sensitivity via neurogenic inflammation, likely leading to localized artemin release that induces cold pain via GFRα3 and TRPM8.

**Significance Statement:** The cellular and molecular mechanisms that generate pain are complex with a diverse array of pain-producing molecules generated during injury that act to sensitize peripheral sensory neurons, thereby inducing pain. Here we identify a specific neuroinflammatory pathway involving the ion channel TRPM8 and the neurotrophin receptor GFRα3 that leads to cold pain, providing select targets for potential therapies for this pain modality.

## INTRODUCTION

Sensations such as touch and pain are initiated by activation of modality-specific sensory receptors on primary afferent nerve endings that respond to mechanical, chemical, or thermal stimuli (Basbaum et al., 2009). After injury, the sensitivity of these receptors is enhanced, thereby lowering thresholds such that pain intensity is increased (hyperalgesia), or stimuli that would normally be innocuous become painful (allodynia). Molecularly, damage to tissues and cells produces a variety of biologically active inflammatory mediators that either directly stimulate pain receptors or enhance their activity indirectly via an assortment of different signal transduction cascades (Woodhams et al., 2017; Ueda, 2021). In addition, robust stimulation of nociceptors induces the antidromic release of neuropeptides, such as calcitonin gene-related peptide (CGRP) and substance P, leading to neurogenic inflammation and further aggravation of the inflammatory and pain states (Woolf and Ma, 2007; Basbaum et al., 2009). Thus, a key factor in the development of novel therapeutic options for acute and chronic pain is a better understanding of the molecules and pathways that lead to peripheral sensitization with injury or disease.

Intriguingly, unlike their effects on heat and mechanical sensitivity, many inflammatory mediators released at the site of injury, such as bradykinin, prostaglandins, and histamine, do not increase cold sensitivity, and may in fact inhibit normal cold responses (Linte et al., 2007; Zhang et al., 2012; Zhang, 2019). This effect involves receptor-mediated inhibition of transient receptor potential melastatin 8 (TRPM8) (McKemy et al., 2002), the principal sensor of environmental cold in mammals that is required for appropriate responses to acute cold and pathological cold pain, thereby diminishing the potential analgesic effects of cooling (Bautista et al., 2007; Colburn et al., 2007; Dhaka et al., 2007; Knowlton et al., 2013). Nonetheless, like other somatosensory modalities, cold is a source of serious discomfort and pain (McKemy, 2018), and a better understanding of the molecules and signaling pathways that lead to cold allodynia is critical to any potential treatments.

The ion channels transient receptor potential vanilloid 1 (TRPV1) and transient receptor potential ankyrin 1 (TRPA1) are expressed in a heterogeneous subpopulation of primary sensory neurons that consists of peptidergic and non-peptidergic C-fiber and Aδ-fiber nociceptors (Caterina et al., 1997; Story et al., 2003; Jordt et al., 2004). Stimulation of these channels results in neuropeptide release which elicits plasma protein extravasation and vasodilatation, collectively referred to as neurogenic inflammation (Woolf and Ma, 2007; Nassini et al., 2014; Matsuda et al., 2019). While TRPV1 is considered a noxious heat receptor (Caterina et al., 2000), TRPA1 was initially identified as a sensor of noxious cold (Story et al., 2003), although this has been under considerable debate in the field after TRPA1 was shown to be a receptor for environmental irritants (Jordt et al., 2004; Caspani et al., 2009). Nonetheless, genetic and pharmacological studies in vivo support a prominent role for TRPA1 in cold pain after injury, consistent with its established function as a key mediator of chronic inflammation (Bautista et al., 2013).

Here, we report that endogenous inflammatory mediators that stimulate neurogenic inflammation via activation of TRPA1 and TRPV1 lead to cold allodynia in a TRPM8-dependent manner. This cold pain phenotype is attenuated by block of both CGRP and substance P signaling, and we find that these neuropeptides can also directly provoke cold allodynia. Intriguingly, CGRP induces allodynia in female but not male mice, and antagonizing CGRP receptors only blocks neurogenic inflammation-induced cold sensation in females, whereas TLR4 inhibition prevents cold allodynia in males. Regardless of which cold pain inducer is used, we find that all require the availability of the neurotrophin artemin and its cognate receptor glial cell line-derived (GDNF) receptor alpha 3 (GFRα3) (Baloh et al., 1998a; Baloh et al., 1998b; Honma et al., 2002). Thus, our study reveals new mediators of cold pain that work via divergent signaling pathways, but all converge on artemin signaling via GFRα3 leading to increased cold sensitivity via TRPM8.

## MATERIALS and METHODS

### Animals

All experiments were approved by the University of Southern California Institutional Animal Care and Use Committee and in compliance with National Institute of Health guidelines. Mice aged from 8 to 14 weeks were used in all experiments and equal numbers of male and female mice were used unless otherwise specified. All mice were housed in a temperature-controlled environment on a 12-hour light and dark cycle with ad libitum access to food and water. All experiments were performed during the light cycle from 9AM to 5PM. Wildtype mice (Strain #000664), TRPM8-null mice (Trpm8^-/-^, Strain # 008198), and TRPA1-null mice (Trpa1^-/-^, Strain #006401) were purchased from Jackson Laboratories and were on the C57Bl/6 background. GFRα3 null mice (Gfrα3^-/-^) (Honma et al., 2002; Lippoldt et al., 2016) were maintained inhouse, and heterozygous mice were bred to obtain wildtype littermate controls (Gfrα3^+/+^).

### Pharmacological Agents

All agents were purchased from the vendors listed in Table 1 and diluted to the desired concentration before being administered via an intraplantar injection in their respective vehicle solutions. All injections were in 20μl except for substance P (35μl). The anti-artemin monoclonal antibody and control IgG isotype antibodies were purchased from R&D Systems, reconstituted in sterile PBS solution, and injected intraperitoneally at a concentration of 10mg/kg (Lippoldt et al., 2016).

**Table 1.**
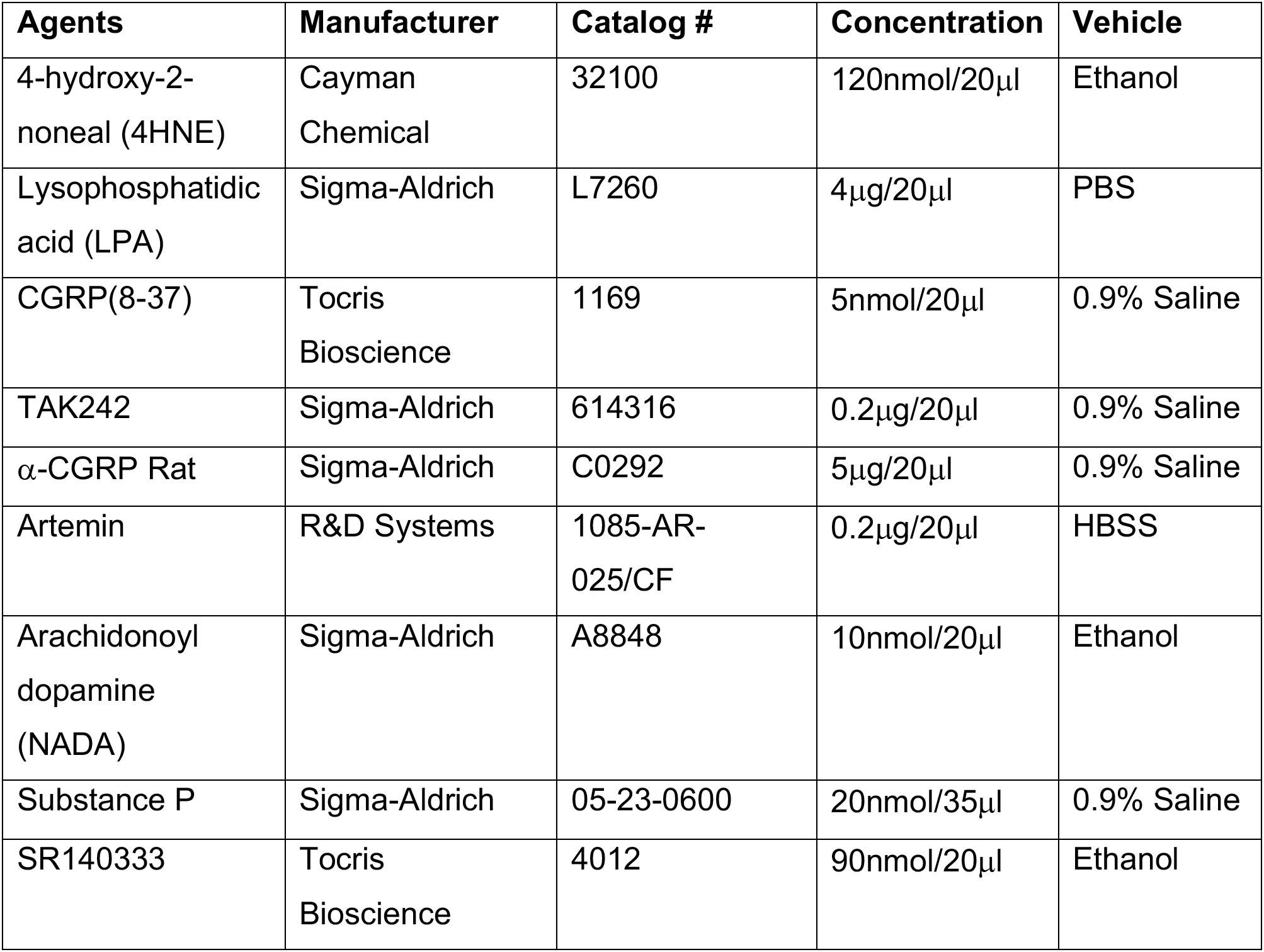

### Cold plantar assay

The cold plantar assay was used to assess the sensitivity to cold stimulation as previously described (Brenner et al., 2014; Ongun et al., 2018; Yamaki et al., 2021). Mice were habituated in plexiglass chambers for 2 hours on a 6mm-thick glass surface maintained at 30°C. A compressed dry ice pellet was applied to the surface of the glass directly under the hind paw to be tested and hind paw withdrawal latencies were recorded. A 30-second cutoff time was used between each measurement to avoid potential tissue damage. Three trials were taken and averaged for each hind paw and baseline measurements were recorded before each experiment. In time-course experiments, mice were returned to home cages after measurements at 4 hours to access food and water and returned to their original chambers for 30 minutes for habituation prior to next measurements.

### Hargreaves assay

The Hargreaves assay was used to assess the sensitivity to heat stimulation (Yamaki et al., 2021). Mice were habituated in plexiglass chambers (IITC) for 2 hours on a glass surface of which temperature was maintained at 32°C. A radiant light source was used as a heat stimulus and focused on the plantar surface of hind paw. Hind paw withdrawal latency was recorded with a cutoff time of 20 seconds between each measurement to avoid potential tissue damage. Three trials were taken and averaged for each hind paw. Baseline measurements were recorded before treatment. Mice were returned to original chamber after treatment and new measurements were taken after 30 or 60 minutes.

### Statistical Analysis

The experimenters were blinded to either genotype of the mice receiving treatments or substances injected and a Power analysis was performed to determine an appropriate sample size. All data was presented as mean and ±SEM and all statistical analysis was performed using Prism Graphpad 9 software as described in the Figure Legends.

## RESULTS

### Endogenous inflammatory mediators induce cold allodynia

The mechanisms whereby inflammatory mediators, produced by tissue injury or other physical insults, lead to cold pain are poorly understood. To address this gap in our knowledge, we tested the effects of established endogenous proalgesic molecules, with distinct receptor mechanisms, and their ability to induce cold sensitivity in mice. First, we tested 4-hydroxy-2-nonenal (4HNE), a reactive aldehyde produced when reactive oxygen species (ROS) peroxidize membrane phospholipids and a known endogenous agonist of the irritant receptor TRPA1 (Benedetti et al., 1980; Trevisani et al., 2007). Using the Cold Plantar assay, a measure of an animal’s sensitivity to an innocuous cold stimulus (Brenner et al., 2014; Ongun et al., 2018), we measured baseline latencies for a hind paw lift upon cold stimulation in an equal number of male and female C57/Bl6 wildtype mice, then performed a unilateral intraplantar hind paw injection of 4HNE (120nmol) or vehicle, with cold sensitivity re-assessed over a 12hr period. 4HNE produced a reduction in withdrawal latencies that was significantly different than baseline by 30mins post-injection, an effect that remained for up to 8hrs (Fig. 1A). The area over the curve (AOC) for each time course showed that 4HNE induced significant cold allodynia compared to vehicle treated animals or the uninjected contralateral hind paw (Fig. 1B). To determine the receptor dependency of 4HNE on cold allodynia, we repeated these assays in TRPA1-null mice (Trpa1^-/-^) (Kwan et al., 2006; Trevisani et al., 2007) finding that the 4HNE-induced reduction in the withdrawal latency observed in wildtype littermates (Trpa1^+/+^) was absent in Trpa1^-/-^ mice (Fig. 1C).

**Figure 1.**
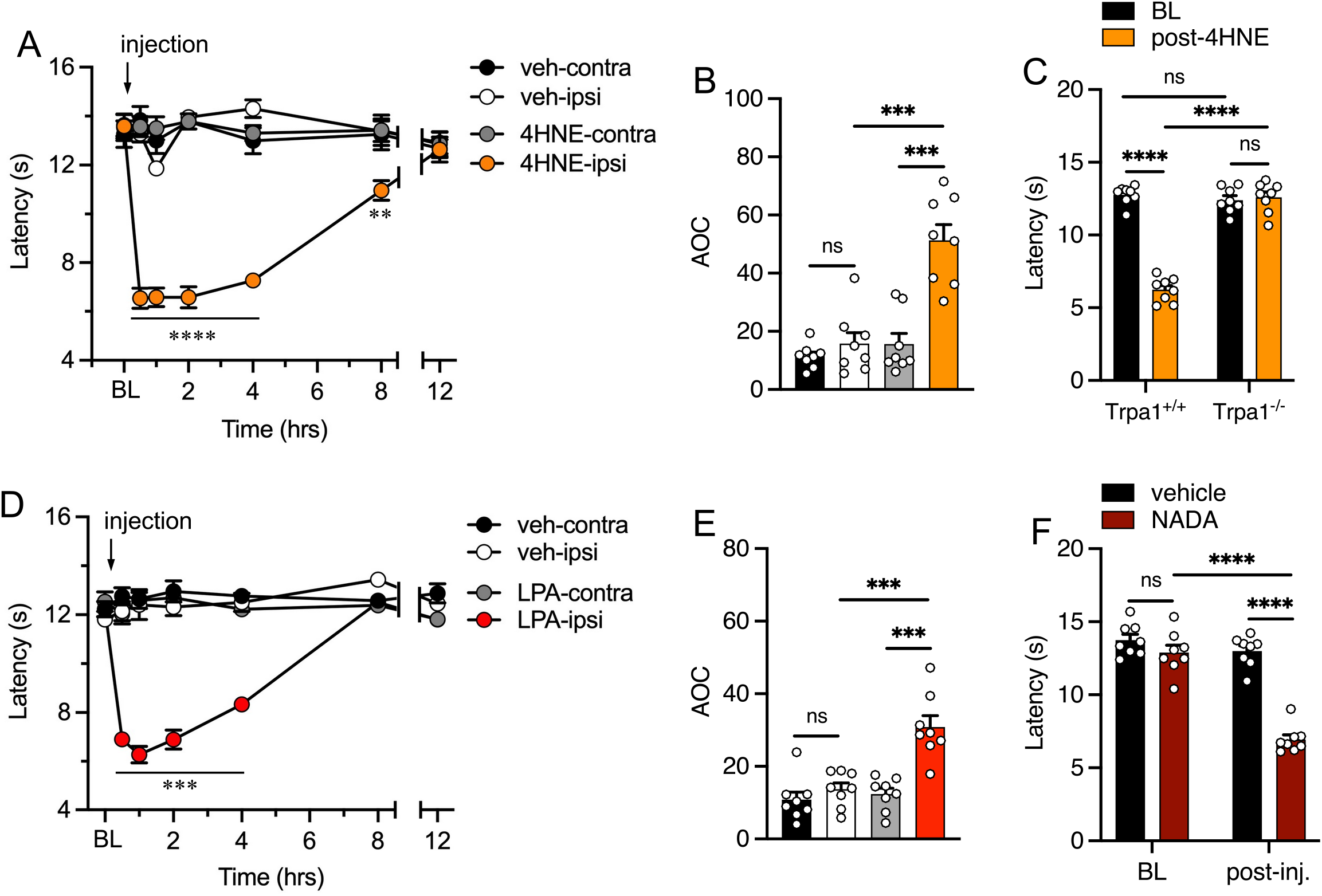
Endogenous inflammatory mediators induce cold allodynia. (**A**) Cold allodynia in mice receiving an intraplantar injection of 4HNE starts at 30min postinjection and lasts up to 8hrs as evidenced by a significantly lower withdrawal latency compared to baseline (BL) measurements or to vehicle injected mice (n=8 for each condition, two-way ANOVA with a Dunnett’s multiple comparisons test for BL versus each time point and Bonferroni’s multiple comparisons test for vehicle versus 4HNE at each time point, ****p<0.0001, **p<0.01). (**B**) Area over the curve (AOC) was measured for each animal over time and averaged between groups, demonstrating that 4HNE produced robust cold allodynia in wildtype mice (paired two-tailed t-test for ipsi versus contra, unpaired t-test with Welch’s correction for vehicle versus 4HNE or LPA, ^ns^p>0.05, ***p<0.001). (**C**) TRPA1-null (Trpa1^-/-^) mice displayed normal cold sensitivity after injection with 4HNE whereas wildtype (Trpa1^+/+^) littermates showed robust cold allodynia (n=8 for each genotype, paired two-tailed t-test for BL versus post-4HNE, unpaired t-test with Welch’s correction for post-4HNE between genotypes, ^ns^p>0.05, ****p<0.0001). Cold sensitivity was similar between genotypes prior to injection (unpaired t-test with Welch’s correction, ^ns^p>0.05). (**D**) Intraplantar LPA also produced robust cold allodynia for at least 4hrs post injection compared to baseline or vehicle (n=8 for each condition, two-way ANOVA with a Dunnett’s multiple comparisons test for BL comparisons and Bonferroni’s for vehicle, ***p<0.001). (**E**) LPA-induced cold allodynia was also significantly different in mice given vehicle or the contralateral hind paw (paired two-tailed t-test for ipsi versus contra, unpaired t-test with Welch’s correction for vehicle versus 4HNE or LPA, ^ns^p>0.05, ***p<0.001). (**F**) Intraplantar NADA induced cold allodynia (n=8, unpaired t-test with Welch’s correction for vehicle versus NADA, paired two-tailed t-test for BL versus post-NADA, ^ns^p>0.05, ****p<0.0001).

Next, we sought to determine if the sensitization induced by 4HNE was a general effect by asking if molecularly and functionally distinct endogenous inflammatory mediators also induce cold allodynia. Specifically, we focused on lysophosphatidic acid (LPA), a phospholipid produced by activated platelets and microglia upon tissue damage (Ma et al., 2010a; Nieto-Posadas et al., 2011). LPA is an agonist of the capsaicin receptor TRPV1 (Huang et al., 2002; Nieto-Posadas et al., 2011) which, like TRPA1, when activated induces neurogenic inflammation and peripheral sensitization (Basbaum et al., 2009). Intriguingly, we found that an intraplantar injection of LPA (4μg) also produced cold allodynia that lasted for several hours (Fig. 1D) and was significantly different compared to vehicle and the contralateral paw (Fig. 1E). To confirm that this effect is not molecule specific, we also tested the endogenous TRPV1 agonist *N*-arachidonoyldopamine (NADA), a putative endocannabinoid that induces robust heat hyperalgesia via activation of TRPV1 (Huang et al., 2002), observing that NADA produced similar effects on cold sensitivity as 4HNE and LPA in wildtype mice (Fig. 1F). Thus, inflammatory mediators produced by tissue insults lead to cold hypersensitivity and, to the best of our knowledge, the effects of these molecules on cold pain, and their associated cellular mechanisms, have not been determined.

While TRPA1 is proposed to be a cold sensor (Story et al., 2003), this has been controversial (Sinica and Vlachova, 2021) and, as we have previously reported (Yamaki et al., 2021), we observed no differences in baseline responses to cold between the two genotypes (Fig. 1C), suggesting that TRPA1 does not mediate cold responses in the cold plantar assay. Furthermore, TRPV1 is a noxious heat sensor and has not been linked to acute cold sensing, suggesting that the effects of LPA or NADA are downstream of TRPV1 channel activation. Thus, it is unlikely that cold allodynia induced by these endogenous molecules is mediated directly by either TRPV1 or TRPA1. The menthol receptor TRPM8 is the predominant cold sensor in mammals and we asked if cold allodynia induced by 4HNE and LPA is dependent on TRPM8 by repeating these assays in TRPM8-null mice (Trpm8^-/-^) (Bautista et al., 2007; Knowlton et al., 2010). To test 4HNE and LPA, both wildtype and Trpm8^-/-^ mice were first tested for cold sensitivity prior to a single intraplantar injection of either 4HNE (Fig. 2A) or LPA (Fig. 2B) as above, then cold withdrawal latencies were re-tested 30mins (for 4HNE) or 60mins (for LPA) later when we observed no differences in cold acuity in Trpm8^-/-^ mice pre-versus post-injection. To ensure that this phenotype was not due to an overall deficit in sensitization in these animals, we repeated these experiments and measured heat hyperalgesia with the Hargreaves assay (Hargreaves et al., 1988), a behavioral paradigm like the cold plantar assay. As predicted, Trpm8^-/-^ mice displayed no differences in heat sensitivity compared to wildtype mice at baseline, whereas 4HNE (Fig. 2D) or LPA (Fig. 2E) induced heat hyperalgesia in TRPM8-nulls that was indistinguishable from that observed in wildtype mice. These results suggest that cold sensitization produced by these inflammatory mediators is downstream of their receptor channels and converge on sensitizing TRPM8.

**Figure 2.**
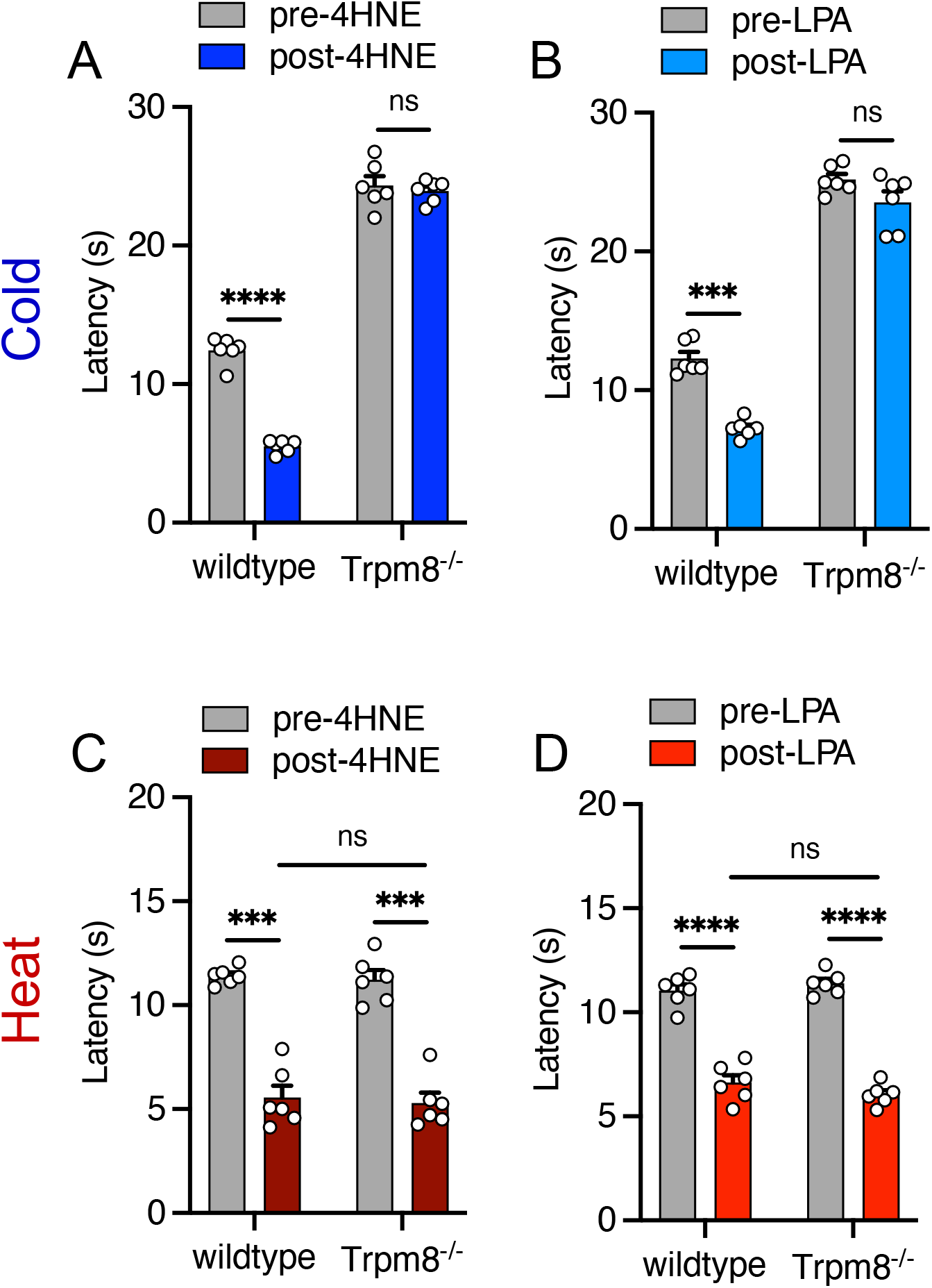
Endogenous inflammatory mediators induced cold allodynia requires TRPM8. An intraplantar injection of either 4HNE (**A**) or LPA (**B**) induced cold allodynia in wildtype but not Trpm8^-/-^ mice (n=6, pre-versus post-injection paired two-tailed t-test, ^ns^p>0.05, ***p<0.001, ****p<0.0001). Conversely, both genotypes exhibited heat hyperalgesia after injection of either 4HNE (**C**) or LPA (**D**), with no differences between wildtype and Trpm8^-/-^ mice either pre- or post-injection (n=6, pre-versus post-injection paired two-tailed t-test, wildtype versus Trpm8^-/-^ post-injection unpaired t-test with Welch’s correction, ^ns^p>0.05, ***p<0.001, ****p<0.0001).

### Substance P and CGRP mediate cold allodynia produced by inflammatory mediators

As stated above, these agents are potent agonists for TRPA1 and TRPV1, nociceptor-expressed non-selective cation channels whose activation leads to neurogenic inflammation and peripheral sensitization to somatosensory stimuli via antidromic release of proinflammatory peptides, such as substance P and CGRP (Basbaum et al., 2009). Thus, to determine how these agents lead to cold pain, we first asked if the cold allodynia they produce was mediated by substance P signaling. To test this, we tested if inhibition of the neurokinin 1 receptor (NK1R) by the peripherally acting, nonpeptide antagonist SR140333 (Oury-Donat et al., 1994) altered cold allodynia induced by intraplantar 4HNE or LPA. First, we performed an intraplantar injection of SR140333 (90nmol) (Teodoro et al., 2013) in an equal number of male and female wildtype mice and tested if basal cold sensitivity was altered 30min post-injection, observing no effect on basal cold sensitivity with NK1R antagonism. Next, these mice were injected with 4HNE (Fig. 3A) and we tested cold responses 30min post-injection, observing a significantly reduced level of cold allodynia compared to mice pretreated with vehicle. However, SR140333 did not completely prevent cold allodynia as there was a slight but significant reduction in the withdrawal latencies compared to baseline and after injection of SR140333. We repeated this assay with LPA, finding that pretreatment with SR140333 reduced LPA-induced cold allodynia compared to those pre-treated with vehicle (Fig. 3B). Unlike 4HNE, there was no difference between the measurements taken after SR140333 injection and those of antagonist plus LPA, whereas there was again a subtle yet significant difference between the latter and baseline responses (Fig. 3B). Next, we asked if substance P can directly induce cold allodynia in mice and if this was also TRPM8-dependent. Wildtype female and male mice (Fig. 3C) were tested for basal cold sensitivity then given either an intraplantar injection of vehicle or substance P (20nmol) then re-tested 30min later when we observed robust cold allodynia, a phenotype that was dependent on TRPM8 as it was absent in male and female Trpm8^-/-^ mice (Fig. 3D). Furthermore, this TRPM8-dependence is cold specific as Trpm8^-/-^ mice of both sexes exhibited heat hyperalgesia with intraplantar substance P (Fig. 3E).

**Figure 3.**
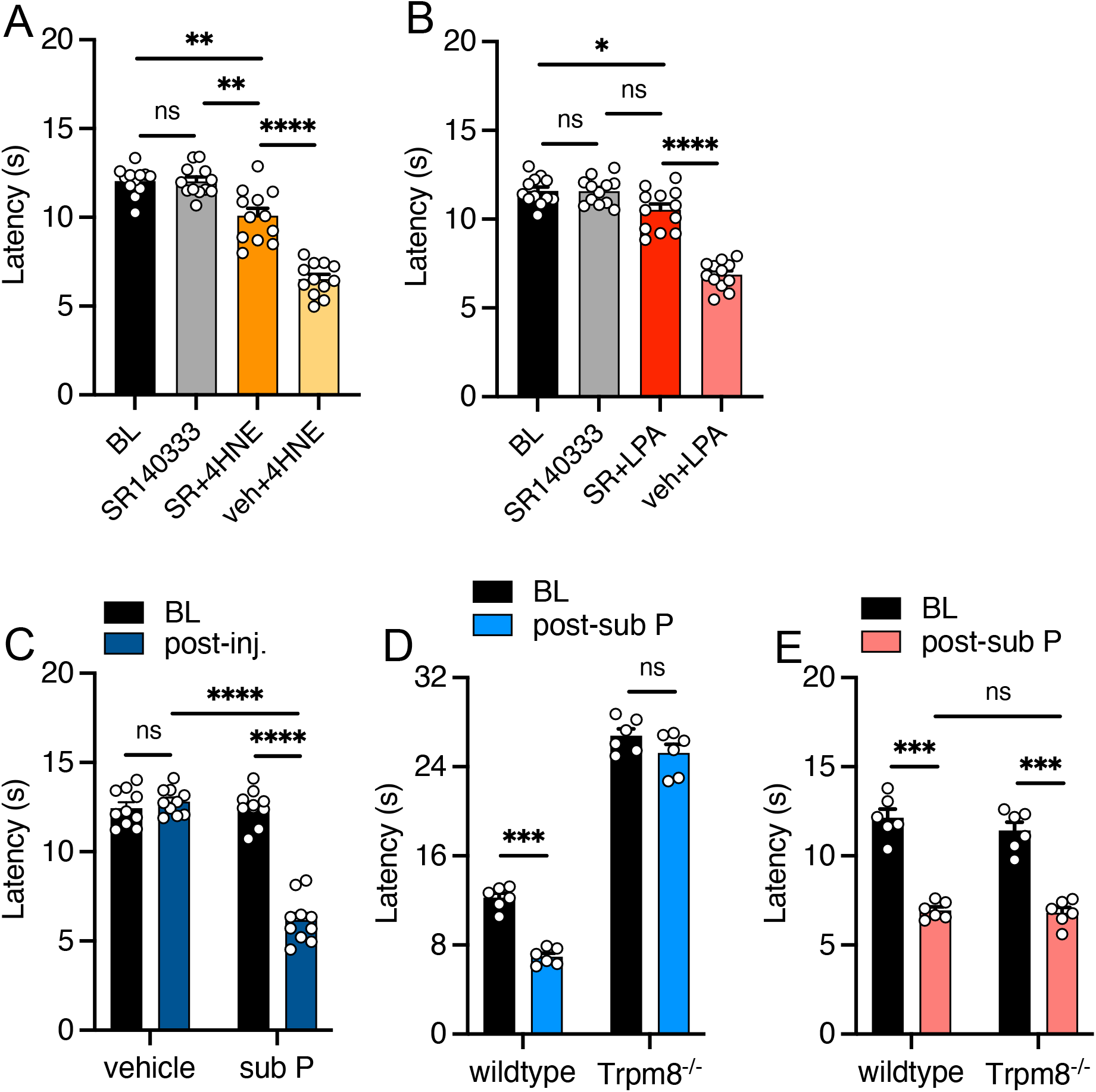
Substance P leads to TRPM8-dependent cold allodynia. Male and female wildtype mice exhibited reduced cold withdrawal latencies after intraplantar injections of 4HNE (**A**) or LPA (**B**) when pretreated with SR140333 (n=12 mice each condition, one-way ANOVA with a Tukey’s multiple comparison test for BL, SR140333, and SR140333(SR)+4HNE or +LPA; unpaired t-test with Welch’s correction for SR140333(SR)+4HNE or +LPA versus veh+4HNE or +LPA; ^ns^p>0.05, *p<0.05, **p<0.01, ****p<0.0001). (**C**) Intraplantar substance P induced cold allodynia in male and female wildtype mice (n=10, paired two-tailed t-test for BL versus post-injection; unpaired t-test with Welch’s correction for vehicle versus substance P; ^ns^p>0.05, ****p<0.0001). Substance P does not induce cold allodynia in Trpm8^-/-^ mice (**D**) but does produce heat hyperalgesia (**E**) in TRPM8-nulls (n=6 for each, paired two-tailed t-test BL versus post-sub P; unpaired t-test with Welch’s correction wildtype versus Trpm8^-/-^, ^ns^p>0.05, ***p<0.001).

Next, we asked if CGRP might also play a role in TRPM8-dependent cold pain. To this end, 30min after baseline cold sensitivity testing, an equal number of male and female wildtype mice received an intraplantar injection of the CGRP receptor antagonist CGRP_8-37_ (5nmol) then re-tested 30min post-injection and we observed no differences in cold sensitivity with CGRP receptor antagonism. We then injected 4HNE as above and observed a modest reduction in the withdrawal latency compared to baseline or after CGRP_8-37_ treatment (Fig. 4A). However, compared to animals injected with vehicle prior to 4HNE (veh+4HNE), mice treated with CGRP_8-37_ prior to 4HNE exhibited significantly less cold allodynia. We similarly tested the effect of CGRP antagonism on LPA-induced cold hypersensitivity, finding that CGRP_8-37_ inhibited cold allodynia induced by this agent compared to baseline and vehicle+LPA treated mice (Fig. 4B). With either agonist, we observed that the latencies in mice treated with the CGRP receptor antagonist displayed substantial variability, prompting us to take a closer look at the data where we detected a clear sexual dimorphism. Specifically, when we increased the number of animals for each sex, we found that CGRP_8-37_ essentially prevented 4HNE-induced cold allodynia in female mice (Fig. 4C). We did observe a subtle, yet significant difference in female mice given CGRP_8-37_ and LPA compared to baseline (p=0.021), but these latencies were not significantly different compared to those taken after CGRP_8-37_ and were significantly different than mice given vehicle prior to LPA (Fig. 4D). Conversely, male mice were unaffected by pretreatment with CGRP_8-37_ prior to 4HNE (Fig. 4E) or LPA (Fig. 4F), demonstrating that antagonizing CGRP receptors inhibits neurogenic cold allodynia in a sex dimorphic manner.

**Figure 4.**
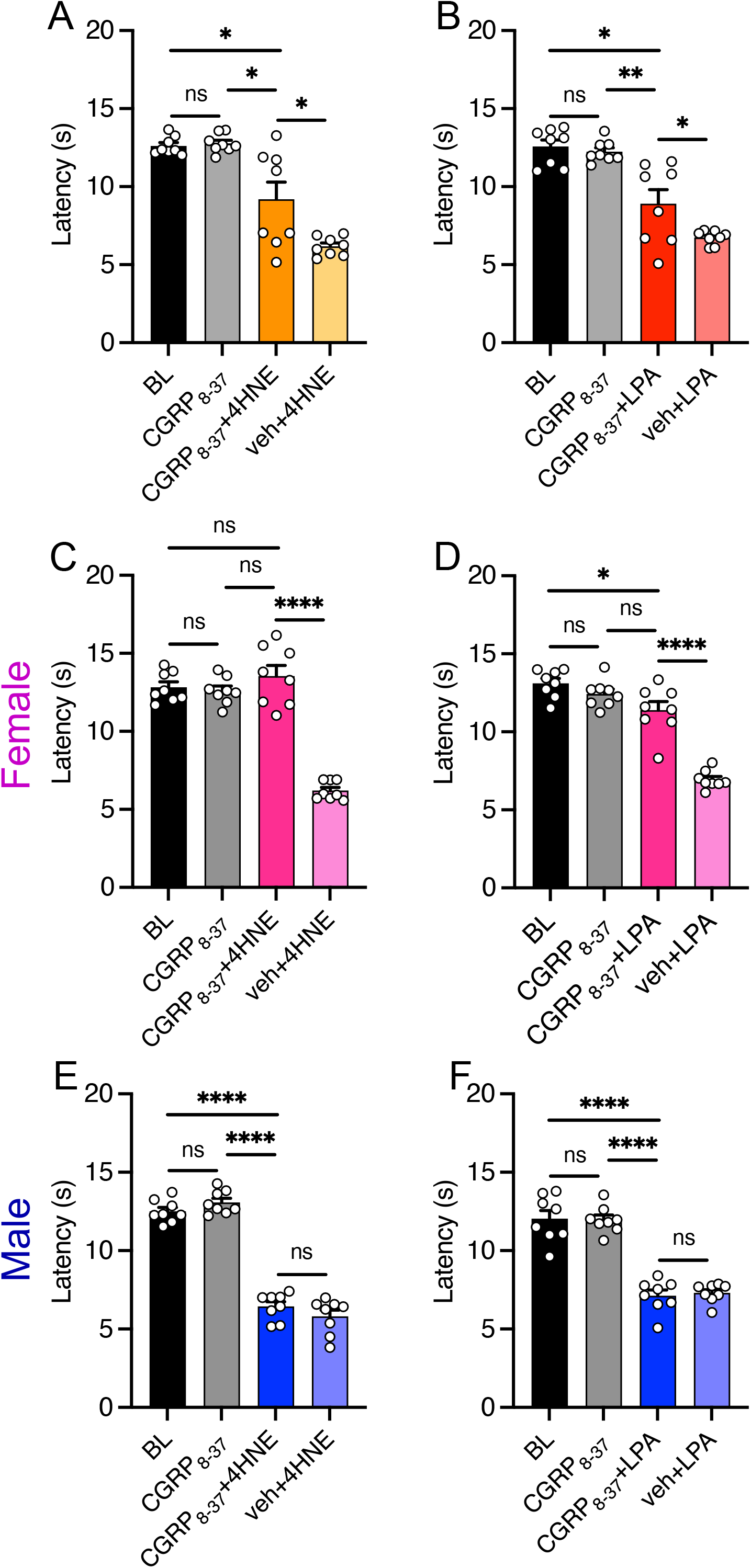
Cold allodynia produced by endogenous inflammatory mediators requires CGRP signaling in female mice. (**A**) Intraplantar injection CGRP_8-37_ in male and female mice did not alter basal cold sensitivity but slightly decreased cold allodynia induced by a subsequent injection of 4HNE (n=8 mice, one-way ANOVA with a Tukey’s multiple comparison test for BL, CGRP_8-37_, and CGRP_8-37_+4HNE; unpaired t-test with Welch’s correction for CGRP_8-37_+4HNE versus veh+4HNE; ^ns^p>0.05, *p<0.05). (**B**) Similarly, CGRP_8-37_ reduced cold allodynia induced by LPA (n=8 mice, one-way ANOVA with a Tukey’s multiple comparison test for BL, CGRP_8-37_, and CGRP_8-37_+LPA; unpaired t-test with Welch’s correction for CGRP_8-37_+LPA versus veh+LPA; ^ns^p>0.05, *p<0.05, **p<0.01). Pretreatment with CGRP_8-37_ essentially prevented cold allodynia induced by either 4HNE (**C**) or LPA (**D**) in female mice. Conversely, CGRP receptor antagonism had no effect on 4HNE- (**E**) or LPA-induced (**F**) cold allodynia in male mice. (n=8 mice for each condition, one-way ANOVA with a Tukey’s multiple comparison test for BL, CGRP_8-37_, and CGRP_8-37_+4HNE or +LPA; unpaired t-test with Welch’s correction for CGRP_8-37_+4HNE or +LPA versus veh+4HNE or +LPA; ^ns^p>0.05, *p<0.05, ****p<0.0001). Data in panels **A** and **B** are included in the sex-specific comparisons in panels **C-F**.

Next, we reasoned that if CGRP receptor antagonism altered cold allodynia, CGRP itself should be cold sensitizing, as well as show sex differences. To directly determine if peripheral CGRP can induce cold allodynia, we performed unilateral hind paw injections of α-CGRP (5μg) or vehicle in both male and female wildtype mice and tested cold sensitivity 30mins post-injection. Consistent with the CGRP receptor antagonist results, α-CGRP produced cold allodynia in female (Fig. 5A) but not male mice (Fig. 5B). Next, we asked if CGRP-induced cold allodynia required TRPM8, finding that the neuropeptide had no effect on cold sensitivity in female Trpm8^-/-^ mice (Fig. 5C). Lastly, to determine if this sexual dimorphism was specific for cold sensing, we repeated the injections of CGRP and assessed heat sensitivity using the Hargreaves’s assay, finding that withdrawal latencies were also reduced in female (Fig. 5D) but not in male wildtype mice (Fig. 5E), and that CGRP did induce heat hyperalgesia in female TRPM8-nulls (Fig. 5F). Thus, peripheral CGRP induces cold allodynia via TRPM8 in a sexually dimorphic manner, consistent with prior reports of female-specific effects of CGRP in peripheral tissues (Paige et al., 2022). Of note, compared to CGRP antagonism, we observed a mildly incomplete inhibition of cold allodynia with NK1R receptor antagonism (Fig. 3) which would not be unexpected as CGRP is likely also altering sensitivity under these conditions. Further, effects of peripheral substance P are not known to show sex differences, and we did not observe any dimorphism when we analyzed our data by sex (p>0.05, multiple unpaired t-tests with Welch’s correction for male versus female mice at each condition in Fig. 3A, B).

**Figure 5.**
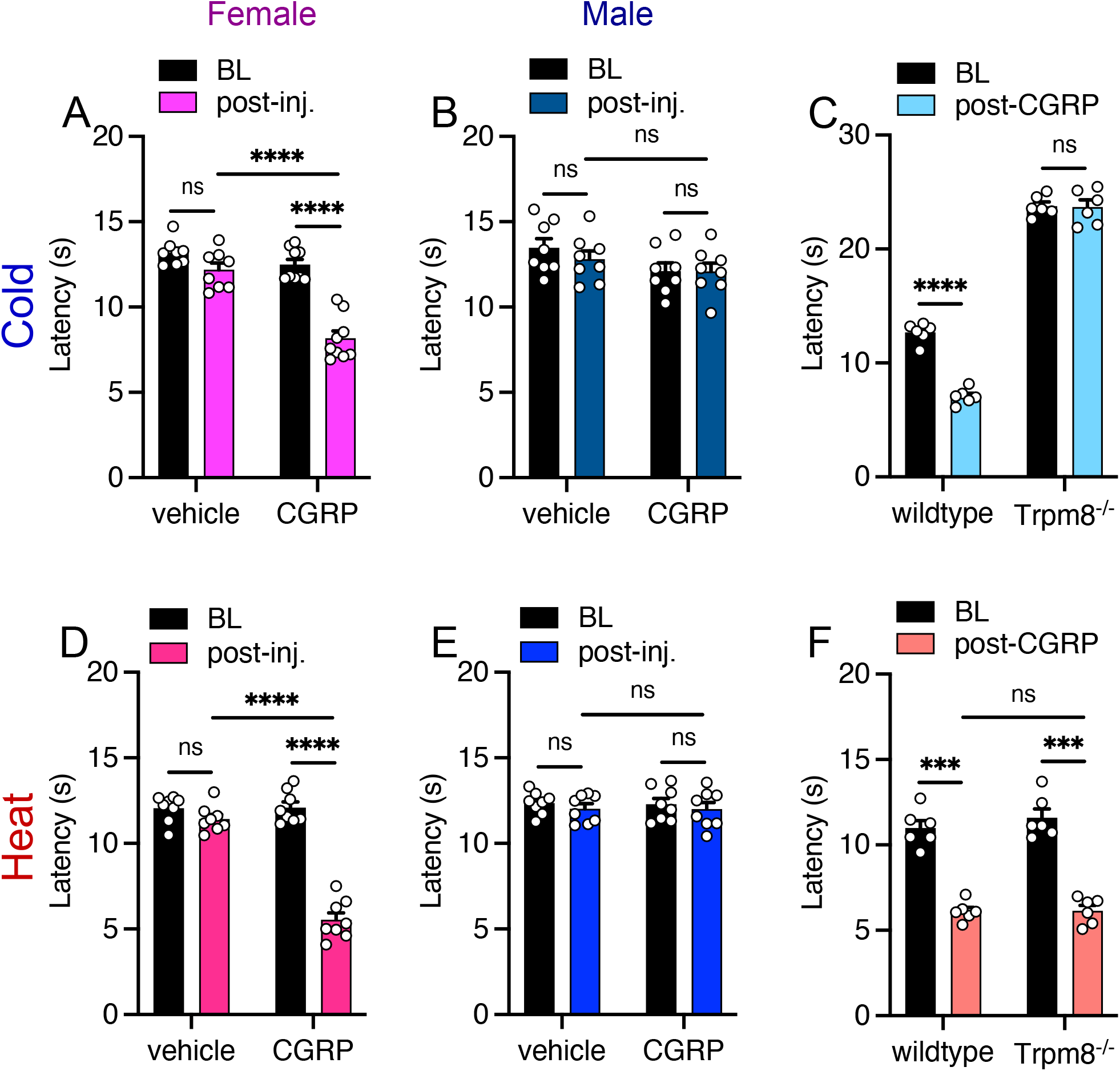
Intraplantar CGRP induces sexually dimorphic cold allodynia. (**A**) Female mice given an intraplantar injection of α-CGRP exhibited robust cold allodynia compared to pre-injection or vehicle injected mice (n=8, paired two-tailed t-test pre-versus post-injection, unpaired t-test with Welch’s correction vehicle versus α-CGRP, ^ns^p>0.05, ****p<0.0001). (**B**) Conversely, α-CGRP did not alter cold sensitivity in male mice (n=8 for each condition, ^ns^p>0.05). (**C**) α-CGRP-induced cold allodynia was absent in Trpm8^-/-^ mice (n=6 for each genotype, paired two-tailed t-test BL versus post-CGRP, ^ns^p>0.05, ****p<0.0001). α-CGRP also induced a similar dimorphic effect of heat sensitivity in female (**D**) and male (**E**) mice (n=8, paired two-tailed t-test pre-versus post-injection, unpaired t-test with Welch’s correction vehicle versus α-CGRP, ^ns^p>0.05, ****p<0.0001). (**F**) α-CGRP-induced heat hyperalgesia was similar in wildtype and Trpm8^-/-^ mice (n=6 for each genotype, paired twotailed t-test BL versus post-CGRP, unpaired t-test with Welch’s correction between genotypes, ^ns^p>0.05, ***p<0.001).

### Inhibition of TLR4 prevents cold allodynia produced by inflammatory mediators in male mice

To further address these sex differences associated with cold allodynia, we sought to determine if other pain signaling pathways known to produce male specific sexual dimorphism also participate in cold allodynia after stimulation by 4HNE or LPA. The Toll-like receptor 4 (TLR4) contributes to microglial-dependent early pain in males, but to a lesser extent in females (Sorge et al., 2011; Woller et al., 2016; Huck et al., 2021), and we asked if inhibiting TLR4 altered cold sensitivity. TAK242 is a small molecule which selectively binds to an intracellular domain of TLR4 and prevents signaling by inhibiting the recruitment of signaling adapter molecules (Matsunaga et al., 2011). Using this antagonist, wildtype female and male mice were tested for baseline cold sensitivity and then given an intraplantar injection of TAK242 (0.2μg), with neither sex exhibiting any alterations in cold sensitivity when re-tested 30min post-injection of TAK242. These animals were then given an injection of 4HNE or LPA as above, with cold sensitivity assessed 30min or 60min later, respectively. Consistent with the male-specific role of TLR4 in pain, we observed no attenuation of 4HNE- (Fig. 6A) or LPA-induced (Fig. 6B) cold allodynia in female mice pretreated with TAK242, whereas this sensitization was completely absent in male mice similarly treated with either 4HNE (Fig. 6C) or LPA (Fig. 6D). Thus, cold allodynia shows intriguing sexual dimorphism with peripheral CGRP and TLR4 signaling required for female and male cold pain, respectively.

**Figure 6.**
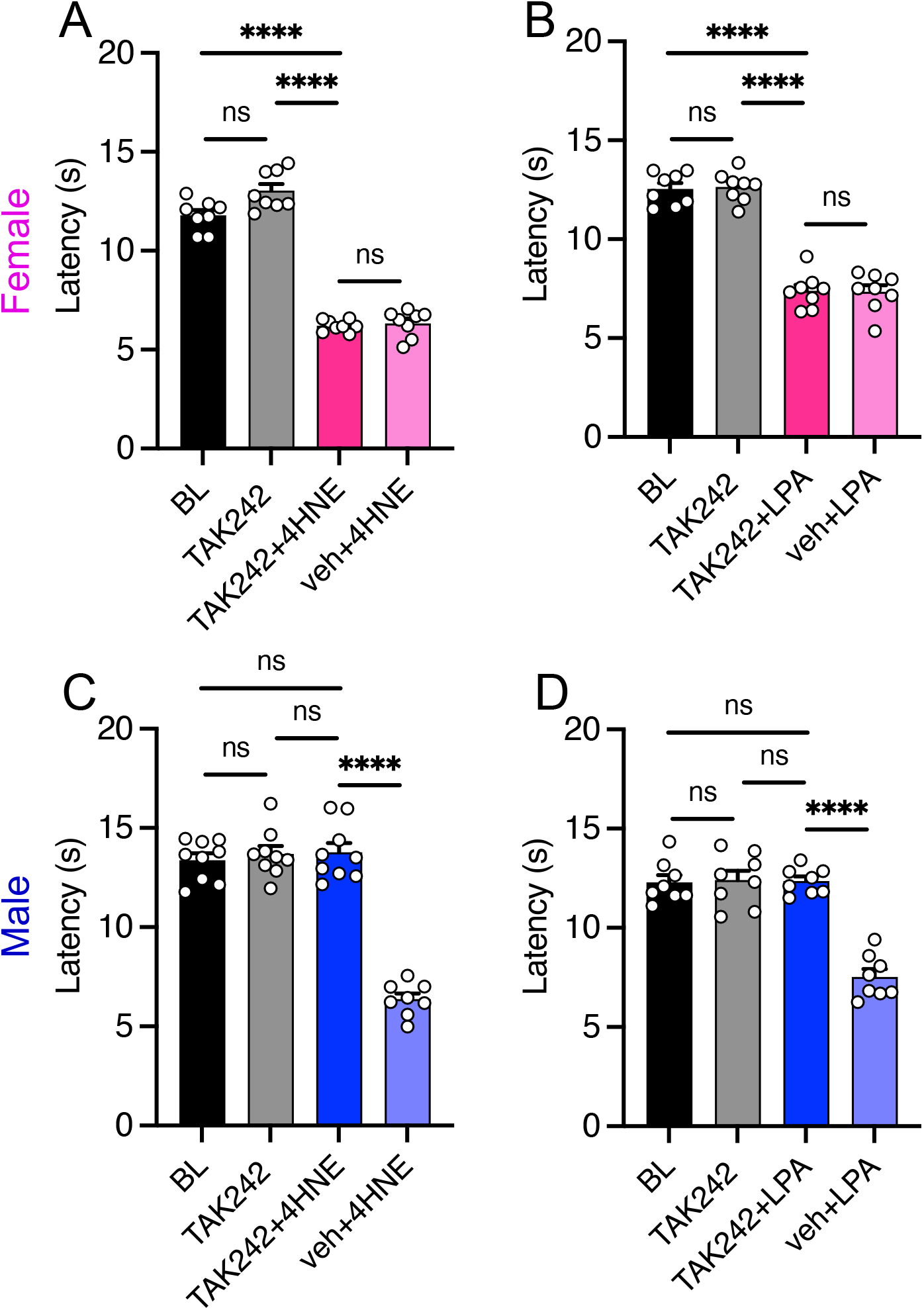
Inhibition of TLR4 prevents cold allodynia produced by endogenous inflammatory mediators in male mice. An intraplantar injection of the TLR4 antagonist TAK242 in female mice did not alter basal cold sensitivity for 4HNE- (**A**) or LPA-induced (**B**) cold allodynia. Conversely, TAK242 prevented cold allodynia induced by 4HNE (**C**) or LPA (**D**) in male mice. (n=8, one-way ANOVA with a Tukey’s multiple comparison test for BL, TAK242, and TAK242+4HNE/LPA; unpaired t-test with Welch’s correction for TAK242+4HNE/LPA versus veh+4HNE/LPA; ^ns^p>0.05, ****p<0.0001).

### Cold allodynia produced by endogenous proalgesics requires the neurotrophin artemin and its receptor GFRα3

When taken together, our results suggest that neurogenic inflammation leads to the release of proalgesic neuropeptides that lead to TRPM8-dependent cold pain. However, how these pathways alter TRPM8 cold signaling is unknown. Previously, we have shown that the neurotrophin artemin sensitizes mice to cold in a manner that is TRPM8-dependent, and that its receptor GFRα3 is required for inflammatory or neuropathic cold allodynia (Lippoldt et al., 2013; Lippoldt et al., 2016). Therefore, we reasoned that neurogenic cold allodynia was similarly dependent on this signaling pathway. To test this we performed intraplantar injections of 4HNE, LPA, CGRP, or substance P in GFRα3-null mice (Gfrα3^-/-^) (Honma et al., 2002; Lippoldt et al., 2016) to determine if this receptor was required for cold allodynia. Both Gfrα3^-/-^ and Gfrα3^+/+^ littermates were tested for basal cold sensitivity and as we have reported previously (Lippoldt et al., 2016), the latencies to lift were similar in both genotypes, as was heat sensitivity (Fig. 7). However, the robust cold allodynia observed in Gfrα3^+/+^ mice after intraplantar injection of either 4HNE (Fig. 7A), LPA (Fig. 7B), CGRP (Fig. 7C), or substance P (Fig. 7D) was absent in Gfrα3^-/-^ animals. Furthermore, while cold allodynia induced by these agents was dependent on GFRα3, heat hyperalgesia remained in Gfrα3^-/-^ mice injected with 4HNE (Fig. 7E), LPA (Fig. 7F), CGRP (Fig. 7G), or substance P (Fig. 7H).

**Figure 7.**
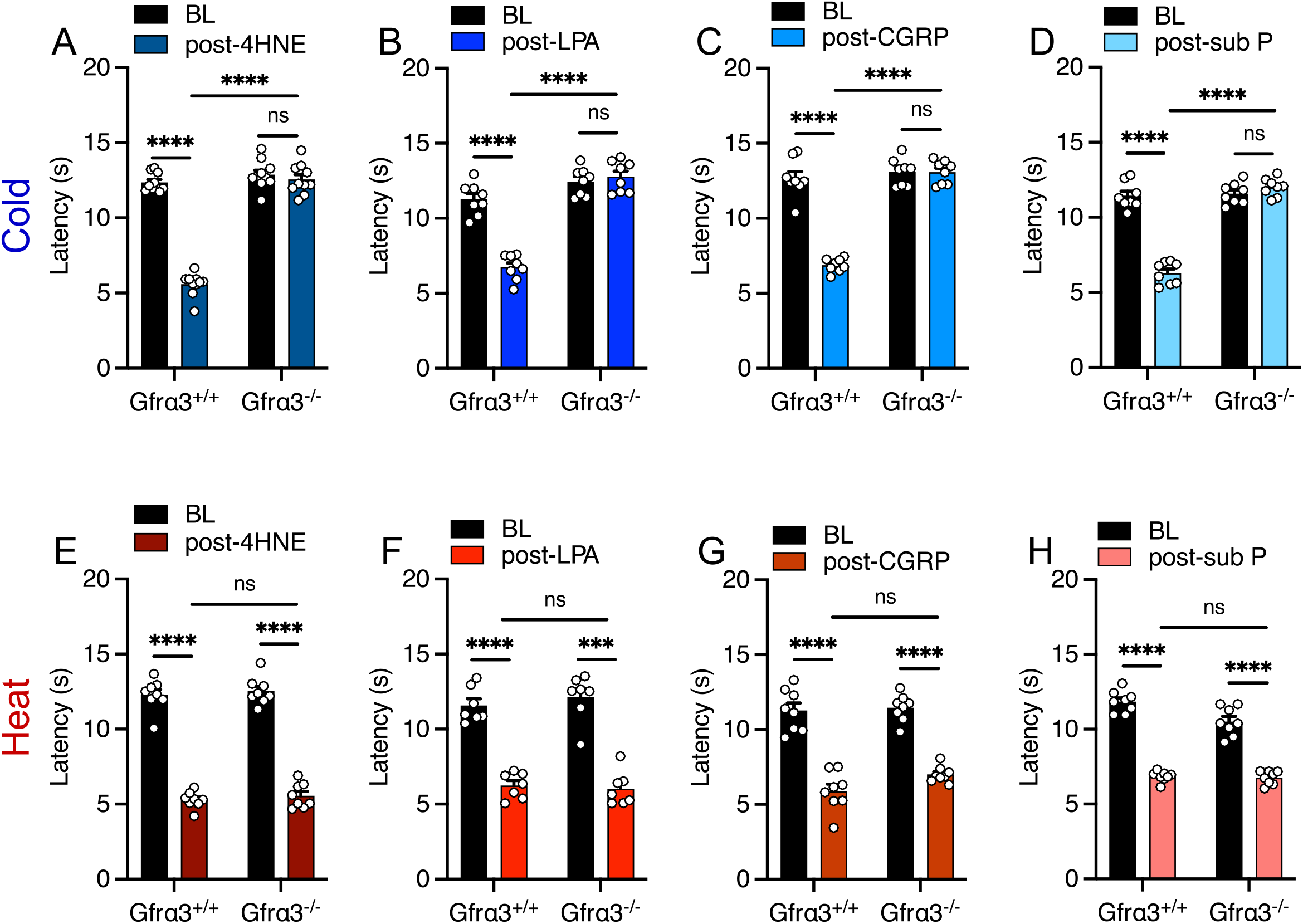
Cold allodynia produced by proalgesics requires GFRα3. Cold allodynia observed in wildtype (Gfrα3^+/+^) mice after injection of either 4HNE (**A**), LPA (**B**), CGRP (**C**), or substance P (**D**) was absent in Gfrα3^-/-^ mice. Conversely, there was no difference in heat hyperalgesia between the two genotypes when injected with either 4HNE (**E**), LPA (**F**), CGRP (**G**), or substance P (**H**) (n=8 each, paired two-tailed t-test BL versus post-injection, unpaired t-test with Welch’s correction post-injection for Gfra3^+/+^ versus Gfrα3^-/-ns^p>0.05, ***p,0.001, ****p<0.0001).

Neutralization of artemin in the periphery selectively resolves both inflammatory and neuropathic cold allodynia (Lippoldt et al., 2016; Jeong et al., 2022). Thus, we asked if this is also true for cold allodynia induced by neuroinflammatory agents injected into the hind paw. Naïve mice were first tested for baseline (BL) cold sensitivity then 4HNE (Fig. 8A), LPA (Fig. 8B), CGRP (Fig. 8C), or substance P (Fig. 8D) were injected then these mice were retested as above, with each agent producing the expected decrease in withdrawal latencies. Immediately after testing, these mice were then given an intraperitoneal (i.p.) injection of either an anti-artemin antibody MAB1085 (10mg/kg) or an equal amount of an isotype control antibody (IgG), with cold sensitivity tested 60min later. For all mice injected with the isotype control, cold allodynia was still present at the time of testing as there was a significant difference in the withdrawal latencies compared to baseline. Furthermore, in the animals treated with LPA (Fig. 8B) or CGRP (Fig. 8C), there were no differences in responses before and after treatment with the control mAb. We did observe a slight yet significant lengthening of the latencies in 4HNE- (p=0.025) and substance P-injected (p=0.001) mice given the control mAb (Fig. 8A, D), but there was still a robust difference compared to baseline. In contrast, we observed an essentially complete amelioration of cold allodynia in mice injected with the anti-artemin mAb (α-ARTN) to all four proalgesics with no significant differences in these withdrawal latencies compared to baseline measurements. In addition, there was a significant difference in cold sensitivity in mice treated with the control versus anti-artemin mAb in all cases. These results indicate that, like both inflammatory and neuropathic cold allodynia, cold sensitization induced by endogenous inflammatory mediators and proalgesic neuropeptides is mediated exclusively by artemin acting on its receptor GFRα3 (Lippoldt et al., 2013; Lippoldt et al., 2016).

**Figure 8.**
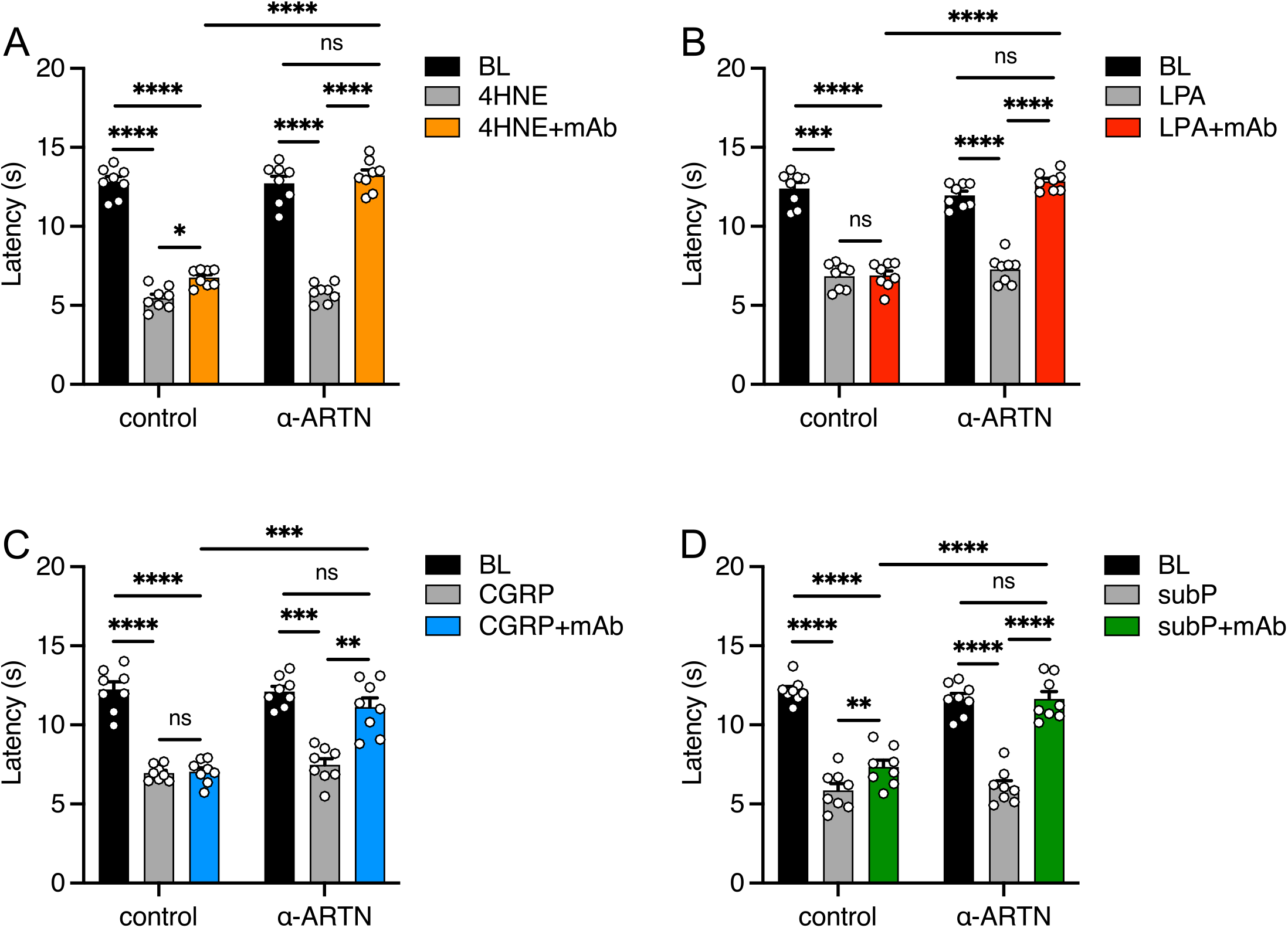
Artemin neutralization ameliorates cold allodynia induced by endogenous proalgesics. Mice treated with control IgG show robust cold allodynia after injection of 4HNE (**A**), LPA (**B**), CGRP (**C**), or substance P (**D**), whereas this cold pain phenotype was ameliorated in mice given an anti-artemin mAb (n=8 for each, one-way ANOVA with Tukey’s multiple comparison tests for BL, proalgesic, and proalgesic+mAb, unpaired t-test with Welch’s correction for control versus anti-artemin mAb comparisons, ^ns^p>0.05, *p<0.05, **p<0.01, ***p<0.001, ****p<0.0001).

### Artemin induces cold allodynia downstream of either CGRP, substance P, or TLR4 signaling

Our data has lent to a potential model whereby activation of nociceptive afferents expressing TRPV1 or TRPA1 triggers CGRP, substance P, and TLR4 signaling pathways which, in turn, prompt the activation of GFRα3, presumably through artemin, and subsequent sensitization of TRPM8 mediated cold sensing (Fig. 9A). This predicts that these signaling pathways are upstream of artemin and, as artemin itself induces TRPM8-dependent cold allodynia (Lippoldt et al., 2013; Lippoldt et al., 2016), we asked if cold allodynia induced by an intraplantar artemin injection could be altered by inhibition of these pathways. To this end, wildtype mice were treated with either CGRP_8-37_ (Fig. 9B), SR140333 (Fig. 9C), or TAK242 (Fig. 9D), or their vehicles, as above, then given an intraplantar injection of artemin 30min later. For all three antagonists, mice treated with artemin showed robust cold allodynia that was not significantly different than mice given their respective vehicles. Thus, inhibition of these signaling pathways does not alter artemin-induced cold allodynia, suggesting that artemin is downstream of the products of tissue injury and neurogenic inflammation.

**Figure 9.**
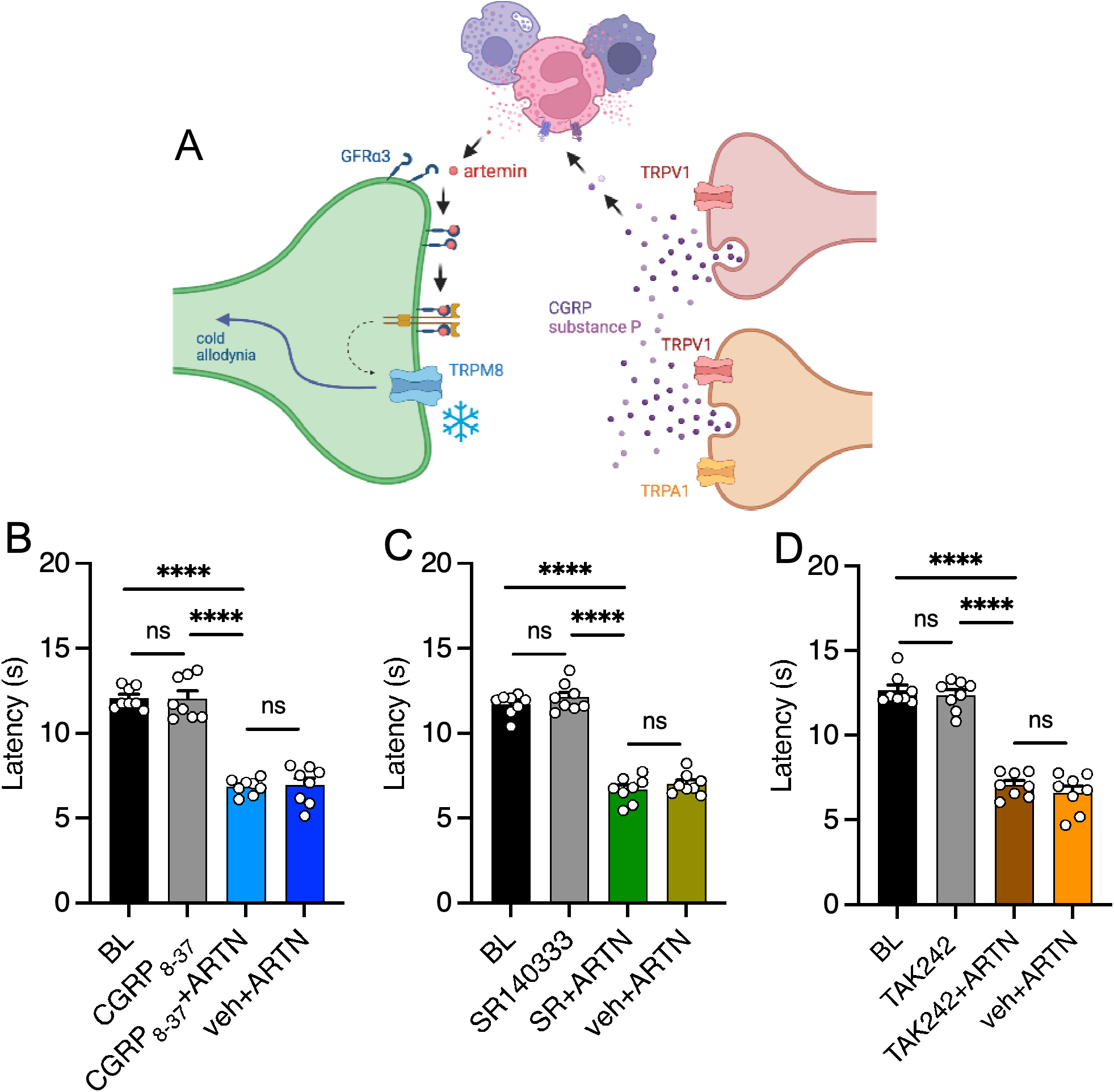
Artemin-induced cold allodynia is insensitive to inhibition of either CGRP, substance P, or TLR4 signaling. (**A**) Model for neurogenically induced cold allodynia (Image generated in BioRender). Wildtype mice pretreated with CGRP_8-37_ (**B**), SR140333 (**C**), or TAK242 (**D**) show robust cold allodynia after an intraplantar injection of artemin (^ns^p>0.05, ****p<0.0001, one-way ANOVA with Tukey’s multiple comparisons test for BL, antagonist, and antagonist+ARTN, unpaired t-test with Welch’s correction for agonist+ARTN versus veh+ARTN).

## DISCUSSION

Here, we show that inflammatory mediators that induce neurogenic inflammation via TRPA1 or TRPV1 lead to cold allodynia involving signaling via CGRP receptors, NK1R, and TLR4. Further, our studies have uncovered an intriguing sex dimorphism in the mechanisms that induce cold hypersensitivity. Regardless of the signaling pathways involved or the sex differences underlying these phenotypes, what is remarkable is that all require the neurotrophin artemin, its receptor GFRα3, and TRPM8. These results are consistent with other pain models, including localized general inflammation, nerve injury, and chemotherapeutic polyneuropathy (Lippoldt et al., 2016), and further highlight the signaling specificity of cold compared to other modalities, such as heat and force in which a plethora of molecules and sensory receptors produce painful sensitization after tissue insults (Basbaum et al., 2009). These findings also point to specific molecular targets that can selectively treat this pain modality without altering sensory processing of either heat or mechanical sensation.

Our results also add to the increasing appreciation of the significant sex differences in pain (Mogil, 2020), in this case with evidence that CGRP and TLR4 mediate cold pain in females and males, respectively. For CGRP, the most well-characterized sex differences occur in migraine-related pain which is significantly more prominent in females compared to males (Ahmad and Rosendale, 2022). Intravenous CGRP triggers headaches in migraineurs (Lassen et al., 2002) and produces migraine-like pain only in female rodents when applied directly to the dura (Avona et al., 2019). In the case of non-migraine pain, surprisingly little is known of potential sex differences of CGRP, likely as most preclinical pain studies have used male mice almost exclusively (Mogil, 2020). However, a recent report found that intrathecally applied CGRP antagonists inhibited hyperalgesic priming in female mice only, and intrathecally applied CGRP caused a long-lasting mechanical hypersensitivity in females, effects that were transient in male mice (Paige et al., 2022).

We show for first-time that peripherally (hind paw) applied CGRP induced thermal hypersensitivity (cold and heat) in female mice only, demonstrating that sex differences also occur in peripheral tissues. Conversely, antagonism of TLR4 prevented neurogenic cold allodynia in males but not females, consistent with some preclinical studies reporting that inhibiting this receptor was only effective at improving pain in males (Sorge et al., 2011; Sorge et al., 2015; Ramachandran et al., 2019), although other studies suggest that the involvement of TLR4 in pain is more complicated (Woller et al., 2016; Huck et al.; Szabo-Pardi et al., 2021). TLR4 is expressed in myeloid-lineage cells both peripherally and centrally, including macrophages, microglia, neurons, astrocytes, and endothelial cells (Ji et al., 2016). Thus, it is unclear where TLR4’s actions lead to sexually dimorphic pain. For example, genetic deletion of TLR4 in sensory neurons expressing Nav1.8 reduced nerve-injury induced mechanical hypersensitivity in female but not male mice (Szabo-Pardi et al., 2021). Further, conditional knockout of TLR4 in microglia reduced chronic allodynia more robustly in male mice in a tibial fracture pain model (Huck et al., 2021). The interaction between CGRP and TLR4 is also interesting and complicated as CGRP modulates inflammatory responses induced by TLR4 activation, whereas TLR4 signaling leads to CGRP release and modulates expression in TLR4-activated macrophages (Gomes et al., 2005; Ma et al., 2010b; Assas et al., 2014; Baliu-Pique et al., 2014; Jia et al., 2021). The TLR4 antagonist used in this study inhibits mechanical allodynia in a postoperative pain model in male rats (females were not tested in this study) when administered peripherally via an intraplantar injection (Xing et al., 2018). Here, we find similar results for neurogenic cold allodynia, suggesting that peripheral TLR4 contributes in a sex-dimorphic manner, but further studies are needed to determine if there are also central effects on cold signaling. TLR4 is likely not co-expressed with TRPM8 neuronally since modulating TLR4 expression in sensory neurons does not induce cold behavioral changes (Szabo-Pardi et al., 2021), which is consistent with our hypothesis of GFRα3 involvement.

Previous studies have shown that substance P release contributes to thermal hyperalgesia (Renback et al., 1999; Massaad et al., 2004; Rogoz et al., 2014), as well as cold nociception in cornea (Li et al., 2019). We find that antagonizing CGRP alone can fully reverse cold allodynia, whereas inhibition of NK1R provided incomplete inhibition. Interactions between CGRP, substance P and other neuropeptides do modulate the effects of substance P in plasma extravasation, and linkages between substance P and TLR4 are also known (Gomes et al., 2005; Rogoz et al., 2014; Schlereth et al., 2016; Huck et al., 2021). Nonetheless, our data is the first to show that intraplantar injection of substance P can induce cold allodynia in mice, regardless of sex. What is striking is that regardless of signaling pathways used herein, each ultimately converges on the release of artemin. Artemin and GFRα3 are known to play important roles in multiple forms of pain, including migraine, osteoarthritis-associated pain and inflammatory bone pain (Shang et al., 2016; Shang et al., 2017; Nencini et al., 2019; Minnema et al., 2022). In addition, there is evidence that artemin can enhance CGRP release from nociceptors (Schmutzler et al., 2009), as well as regulate TRP channels’ expression level (Elitt et al., 2008; Ikeda-Miyagawa et al., 2015). Why artemin has such a specific role in cold nociception, compared to redundant mechanisms in heat and force nociception, remains unknown and requires further analysis.

Lastly, one of the key findings of this study is that it supports the premise that TRPA1 is not a molecular sensor of cold but serves an important role in amplifying cold pain associated with injury as it does for other modalities (Bautista et al., 2013). Since their clonings, TRPM8 and TRPA1 have been considered the primary molecular detectors of cold in the peripheral nervous system, with the premise that due to their temperature activation thresholds in vitro, TRPM8 mediated innocuous cool and TRPA1 noxious cold (McKemy et al., 2002; Story et al., 2003). While the role of TRPM8 in cold is largely incontrovertible, TRPA1 as a molecular sensor of cold has been an intense topic of debate in the field after it was shown to be a receptor for pungent and inflammatory molecules (Bandell et al., 2004; Jordt et al., 2004; Trevisani et al., 2007). In vivo, the strongest evidence for TRPA1 mediating cold has come from studies showing that inflammatory or neuropathic cold pain are lessened in animals with reduced TRPA1 expression or treated with antagonists (Obata et al., 2005; da Costa et al., 2010; del Camino et al., 2010; Palkar et al., 2015). Further, in some cases cold sensitivity is increased when animals are treated peripherally with TRPA1 agonists, as we show here (del Camino et al., 2010; Honda et al., 2014). These convincing data demonstrated that TRPA1 is involved in cold pain in these pathological settings leading to the reasonable supposition that it is a cold sensor in vivo. However, our findings provide a plausible explanation for the in vivo results described above in that activation of TRPA1 and TRPA1-afferents do not directly send cold signals centrally but lead to neurogenic inflammation that promotes sensitization of TRPM8 cold signaling. This premise is further supported by the corollary findings that activation of TRPV1 leads to an identical cold phenotype, including directly inducing cold allodynia, the dependence on CGRP, substance P, and TLR4 signaling, and the dependence on artemin and GFRα3 for this cold pain phenotype.

## Acknowledgments

We thank members of the McKemy Lab for their encouragement and guidance. This study was supported by US National Institutes of Health grant R01 NS106888 (D.D.M.).

